# Identification of a *Nidovirales* Orf1a N7-guanine cap Methyltransferase signature-sequence as a genetic marker of large genome *Tobaniviridae*

**DOI:** 10.1101/639369

**Authors:** François Ferron, Humberto Julio Debat, Etienne Decroly, Bruno Canard

## Abstract

Members of the *Nidovirales* order have (+)RNA genomes amongst the largest in size in the RNA virus world. Expression of their genes is promoted through reading of genomic RNA and mRNA transcripts by the ribosome of the infected cell. The 5’-end of these RNAs is supposedly protected by an RNA-cap structure (m7GpppNm) whose most synthesis steps remain elusive. In Eukaryotes, the RNA-cap structure is methylated by RNA methyltransferases (MTases) at the RNA-cap N7-guanine position as well as the 2’-O methyl position of the first transcribed nucleotide. In *Coronaviridae*, two separate enzymes (nsp14 and nsp16) perform the N7-guanine and the 2’-OH methylation, respectively. One salient feature of the *Nidovirales* N7-guanine MTase nsp14 is that it is the only example of non-Rossman fold viral MTase known so far. Conversely, all other *Nidovirales* nsp16-like MTases have a canonical Rossman fold. Many *Nidovirales* members lack either any RNA MTase signature sequence (e.g., *Arteriviridae*), or lack a N7-guanine MTase signature sequence (e.g., *Tobaniviridae*, *Euroniviridae*, *Roniviridae*, *Medioniviridae*). Both nsp14-and nsp16-like enzyme genes are usually located in Orf1b encoding for the replication machinery. Here, we report the discovery of a putative Rossman fold RNA MTase in the Orf1a of ten *Tobaniviridae* members. Multiple sequence alignments and structural analyses identify this novel gene as a typical RNA-cap N7-guanine MTase with substrate specificity and active-site organization similar to the canonical eukaryotic RNA-cap N7-guanine MTase.

## Introduction

Up to recently, four viral families, *Arteriviridae*, *Coronaviridae*, *Roniviridae*, and *Mesoniviridae* constituted the *Nidovirales* order. The *Coronaviridae* family was further divided in two subfamilies, *Coronavirinae* and *Torovirinae.* The *Torovirinae* family contained two genus *Torovirus* and *Bafinivirus*, plus few unassigned species. Recently, the ICTV proposed a novel classification into the *Nidovirales* order, with the creation of nine families [1]. In this novel ICTV proposal, ratified on February 2019, *Coronaviridae* do not encompass both *Coronavirinae* and *Toronavirinae* subfamilies. Rather, the new suborder *Tornidovirinae* (including toroviruses and bafiniviruses) contains a novel family, *Tobaniviridae*, at the same level as the *Coronaviridae*, which is now within suborder *Cornidovirineae*. This new taxonomy is used here, as depicted in Fig. 1.

**Figure 1.**
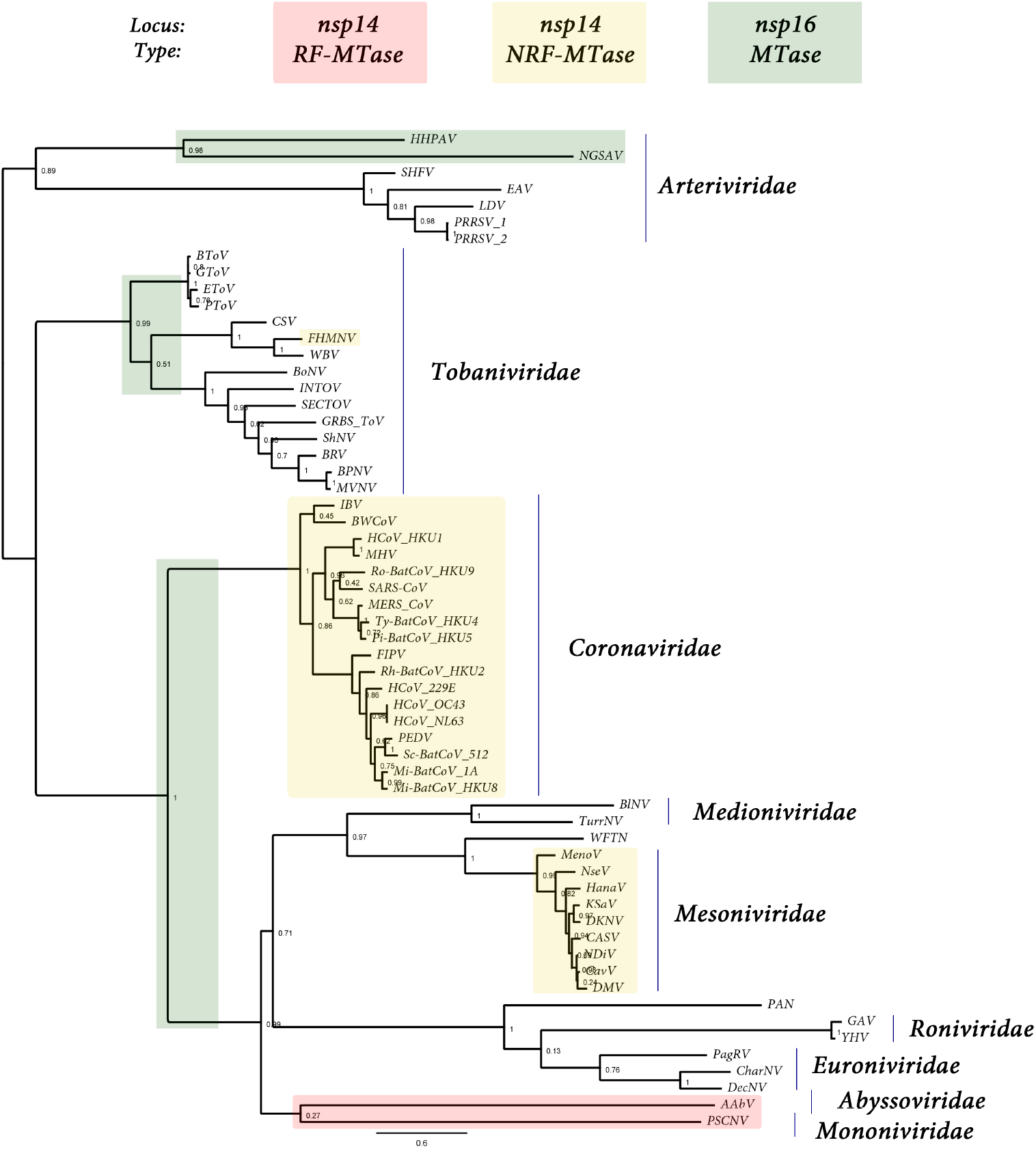
Presence, type, and loci of signature-sequences of *Nidovirales* RNA MTases : RF MT of unknown specificity (light red), non-RF « nsp14-like » MT (yellow), and RF « nsp16-like » MT (green). Both light red and yellow MTases map to the nsp14 C-ter locus, ie., immediately downstrean the ExoN domain, while « nsp16-like » MT map to the nsp16 locus, at the end of Orf1b. The tree was made based on MAFFT v7.427 multiple sequence alignment with BLOSUM62 scoring matrix and G-INS-i iterative refinement method. The alignments were used as input for maximum likelihood trees generated with the FasTtree v2.1.5 software (best-fit model = JTT-Jones-Taylor-Thorton with single rate of evolution for each site = CAT). Local support values were computed using the Shimodaira-Hasegawa test (SH) with 1,000 replicates. Numbers at the nodes represent FasTtree support values and scale var substitutions per site. The tree included two novel Ronivirus-like and Mesonivirus-like genome sequences: Western Flower Thrips Mesonivirus, and Palaemon Nidovirus, respectively (WFTV, PAN, unpublished). When present in the NCBI viral genomes database in the 92 *Nidovirales* complete genomes repository (https://www.ncbi.nlm.nih.gov/genomes/GenomesGroup.cgi?taxid=76804), no accession number is indicated. When an accession number is given in parenthesis, it is referring to the GenBank accession number (https://www.ncbi.nlm.nih.gov/genbank/). From top to bottom of the figure: *Arteriviridae:* HHPAV: Hainan hebius popei arterivirus (MG600021); NGSAV: Nanhai ghost shark arterivirus (MG600024); SHFV : simian hemorrhagic fever virus; EAV: Equine arteritis virus; LDV, Lactate elevating virus; PRRSV-1 and 2, porcine reproductive and respiratory syndrome virus. *Tobaniviridae:* BToV: Breda virus; GToV: Goat torovirus; EToV: Berne virus (CAA36747); PToV: Porcine torovirus; CSV: Chinook salmon bafinivirus; FHMNV: Fathead Minnow nidovirus; WBV: white bream virus; BoNV: Bovine nidovirus TCH5; INToV: Xinzhou nido-like virus 6; SECToV; Xinzhou toro-like virus 1; GRBS-ToV: Guangdong red banded snake torovirus (MG600030); ShNV: shingleback nidovirus 1 (KX184715); BPNV: Ball python nidovirus 1; MVNV: Morelia viridis nidovirus; BRV: Bellinger River virus (MF685025); *Coronaviridae:* IBV: Infectious Bronchitis Virus; BWCoV: Beluga Whale coronavirus SW1; HCoV_HKU1: Human coronavirus HK1; MHV: Mouse hepatitis virus; BatCoV_HKU9: Rousettus bat Coronavirus HKU9; SARS-CoV: Severe acute respiratory syndrome coronavirus; MERS_CoV: middle-east respiratory syndrome coronavirus; Ty-BatCoV_HKU4: Tylonycteris bat coronavirus HKU4; Pi-BatCoV_HKU5: pipistrellus bat coronavirus HKU5; FIPV: Feline infectious peritonitis virus; Rh-BatCoV-HKU2: Rhinolophus bat coronavirus HKU2; HCoV_229E: Human coronavirus 229E; HCoV_OC43: Human coronavirus OC43 (YP_003766); HCoV_NL63: Human coronavirus NL63; PEDV: Porcine epidemic diarrhea virus; Sc-BatCoV_512: Scotophilus bat coronavirus 512; Mi-BatCoV-1A: bat coronavirus 1A; Mi-BatCoV_HKU8: Miniopterus bat coronavirus HKU8. *Medioniviridae:* BlNV: Botrylloides leachii nidovirus; TurrNV: Turrinivirus 1; WFTV: Western Flower Thrips virus. *Mesoniviridae*: MenoV: Meno virus; NseV: Nse virus; HanaV: Hana virus; KSaV: Karang sari virus; DKNV: Dak Nong virus; CASV: Casuarina virus; NDiV: Nam Dinh virus; CavV: Cavally virus; DMV: Dianke mesonivirus. *Roniviridae:* PAN: Palaemon nidovirus; GAV: Gill-associated virus; YHV: Yellow head virus (EU487200). *Euroniviridae*: PagRV: Paguronivirus 1; CharNV: Charybnivirus 1; DecNV: Decronivirus 1. *Abyssoviridae*: AAbV: Aplysia abyssovirus 1 *Mononiviridae*: PSCNV: Planidovirus 1

*Nidovirales* have attracted much attention in the past decades for being causative agents of serious diseases in humans and animals. In humans, Severe Acute Respiratory Syndrome (SARS) and the Middle-East Respiratory Syndrome (MERS) coronaviruses (CoVs) have been associated with high fatality/case ratios of ~10 % ~35 %, respectively (reviewed in [2]). In animals, pathogens of significant economical and societal impact include Equine Arteritis virus (EAV, Alphaarterivirus equid) and Porcine Reproductive Respiratory Syndrome virus (PRSSV 1-2, Betaarterivirus suid 1-2).

Two salient features of these viruses are, amongst others, their complex and still obscure replication/transcription mechanism as well as their broad genome-size range. Genomes of *Arteriviridae* (12.7 −15.7 kb in length), *Mesoniviridae* (~20 kb), *Medioniviridae* (20.2 – 25 kb), *Euroniviridae* (~24.5 kb), *Roniviridae* (~26 kb) *Tobaniviridae* and *Coronaviridae* (~27-32 kb), *Abyssoviridae* (35.9 kb), and *Mononiviridae* (41.1 kb) members, share a similar genomic organisation generating viral mRNAs from a fixed genomic location, the ‘nest’, which has inspired the family name (*Nido* : latin for ‘nest’) [3,4]. *Nidovirales* genomic and mRNAs are presumably capped, although direct demonstration of the presence of an RNA cap structure is still missing for many of these *Nidovirales* members.

In Eukaryotes, the RNA cap structure is thought to be synthesized post-transcriptionally [5]. Most of (+)RNA viruses supposedly follow a conventional RNA capping pathway [6] in which nascent viral 5’-triphosphate genomic (or mRNAs) is thought to be processed through three enzymatic reactions to yield an RNA cap whose structure is indistinguishable from that of cellular mRNAs. The capping pathway implies that the 5’-triphosphate RNA is hydrolyzed into 5’-diphosphate by an RNA triphosphatase, a GMP residue (the « cap », coming from GTP) is next covalently transferred to the 5’-diphosphate RNA in the 5’ to 5’ orientation by a guanlylyltransferase (GTase), releasing inorganic pyrophosphate. Both the cap and the first transcribed nucleotide are then methylated at the N7-guanine (mGpppN-RNA, so-called Cap0 structure) and the 2’-oxygen position (mGpppNm-RNA, Cap1 structure) by one or two RNA methyltransferases (MTases), respectively. The MTase uses S-Adenosyl-Methionine (SAM) as the methyl donor and releases S-Adenosyl-homocysteine (SAH).

There is no high-resolution structural analysis available of *Nidovirales* RNA caps. The evidence for their presence comes from immunological detection in Torovirus genomic and mRNAs [7], and co-migration analysis in low-resolution chromatographic techniques for MHV [8] and Simian Hemorragic fever virus [9]. In other instances, *Nidovirales* cap structures supposedly conform to the canonical eukaryotic RNA capping pathway because enzymes usually deemed necessary to perform RNA capping are present in cognate genomes (reviewed in [6]). Bioinformatic analyses have been used to detect signature-sequences typically associated with RNA capping/modifying enzymes *i.e* guanlylyltransferase (GTase), polymerase, helicase, nuclease, and methyltransferase (MTase). These signature-sequences have served in many instances to isolate corresponding enzymes endowed with the predicted activity. In *Nidovirales*, this approach has been pioneered for Coronavirus enzymes, yet regarding their RNA capping pathway, it only succeeded in predicting 2’OMTase, while a second non-canonical RNA-cap N7-guanine MTase was later structurally and functionally characterized [4,10–14]. Until then, viral MTases matched a specific structural architecture, the Rossmann fold (RF), which possess a unique interaction geometry that represents a fingerprint of common ancestry. The discovery of a non-Rossmann fold (NRF) MTase unveiled the limitation of annotating “dark proteome” data [15], lacking structural counterparts. In *Nidovirales*, enzyme activities have been experimentally evidenced only for coronaviruses and roniviruses [16]. The putative first step of the *Nidovirales* RNA capping pathway, involving a 5’-triphosphatase, is uncertain: such activity exists in the Coronavirus helicase (nsp13, [17]), but its involvement in RNA capping has not been directly demonstrated. The guanylyltransferase also remains unknown: A nidovirus RdRp-associated nucleotidyltransferase (NiRAN) activity was found to be associated to the N-terminus of the arterivirus nsp9 RdRp [18], but its role in arterivirus RNA synthesis, if any, remains to be determined.

Despite the conservation of MTases amongst large *Nidovirales* genomes, their distribution, type, and the function of MTases in the *Nidovirales* order is not homogenous (summarized in Table 1, see text). Arterivirus or arterivirus-like genomes are not known to carry any obvious MTase signature sequence; Mesonivirus and Ronivirus genomes code for one 2’-O RF-MTase (nsp16-like) signature-sequence; The *Tobaniviridae* (*e.g*., toroviruses and bafiniviruses, see Fig. 1) carry a single 2’-O RF-MTase (nsp16-like) signature sequence, whereas *Coronaviridae* carry both a N7-guanine non-RF-MTase (NRF-MTase, nsp14-like) and a 2’-O RF-MTase (nsp16-like) signature-sequence. A single deviation for this MTase-based grouping is that of Fathead minnow nidovirus 1 (*Tobaniviridae*) which carries both MTases (see below) as in the case of *Coronaviridae*.

**Table 1.**
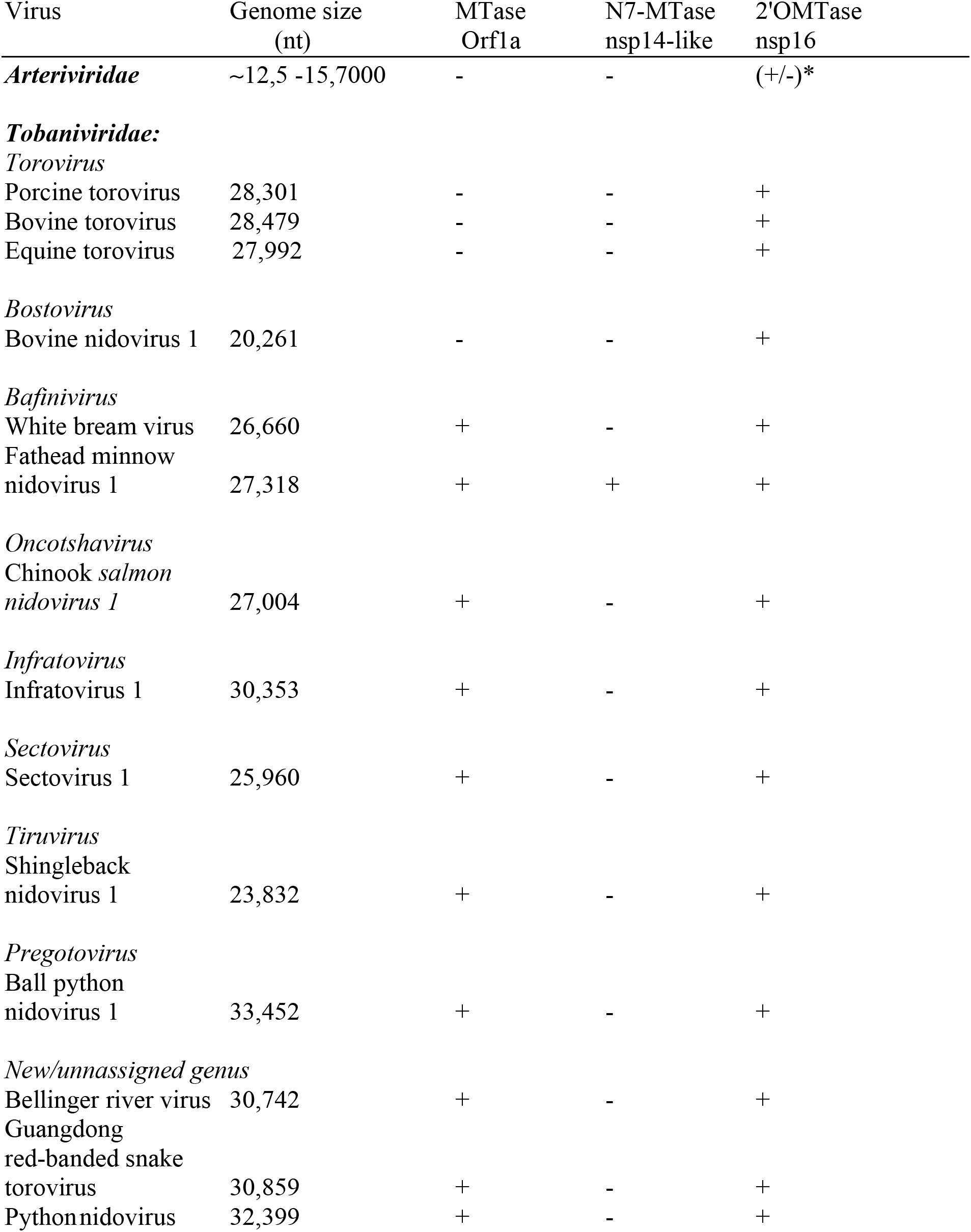
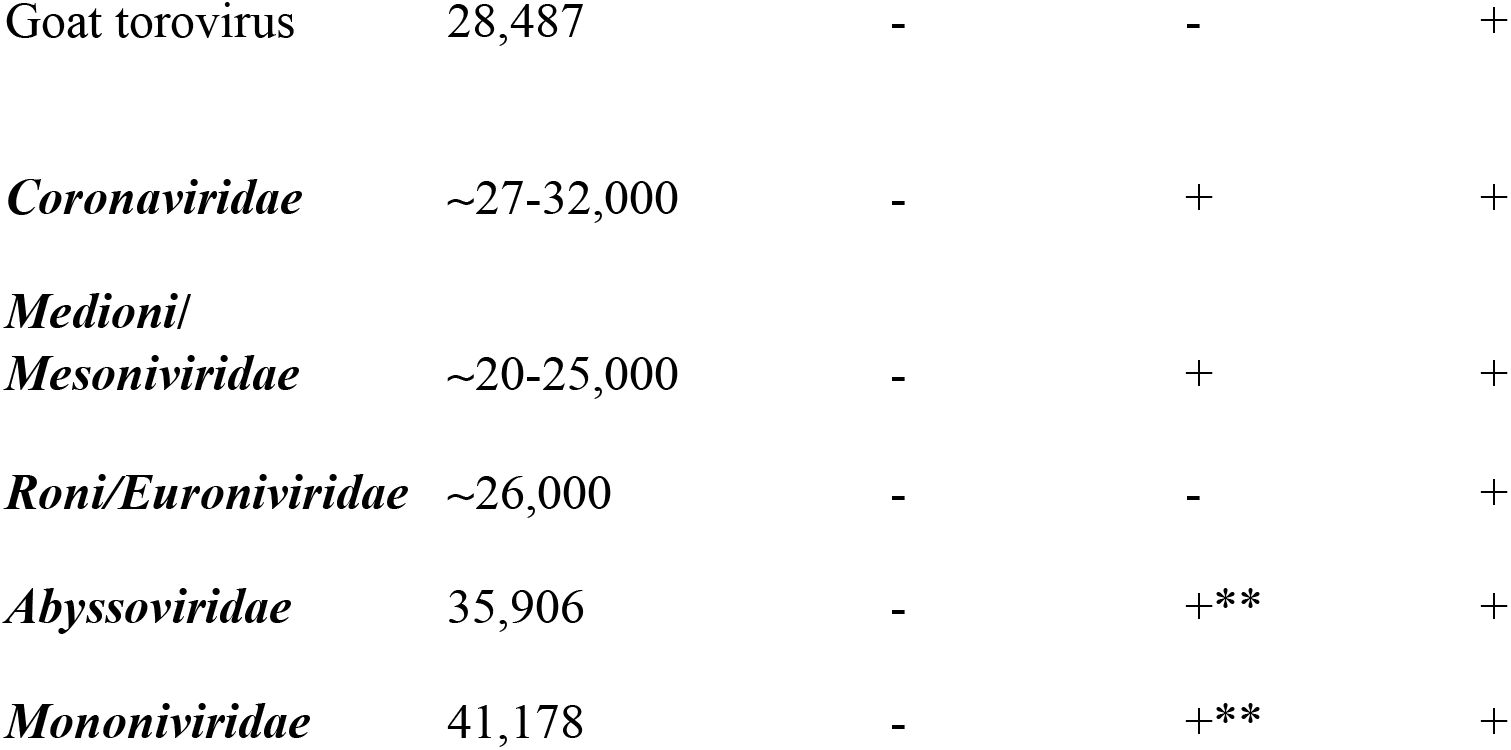
Distribution and putative substrate specificities of MTases in the four families of the *Nidovirales* Order. Only the *Tobaniviridae* is expanded and genera indicated in italic plus unassigned viruses (see Fig. 1). *: Arteriviruses do not usually carry any MTase, except the two newly identified Hainan Hebius Popei arterivirus and Nanhai gost shark arterivirus (see text).**: although in the nsp14 locus immediately downstream the ExoN gene, the MTase has a Rossmann fold. Virus species names are indicated and genome size corresponds to exemplar virus sequence used in this study.

The Coronavirus nsp14 is a peculiar, bi-functional enzyme combining in N-terminus a 3’-to-5’ exonuclease domain (ExoN) fused to the N7-guanine NRF-MTase domain in C-terminus [4,11,12,19,20]. First, it is the only known example of a viral SAM-dependent RNA MTase (and more generally, a unique example of an RNA-Cap 0 N7-guanine MTase) that does not have a Rossman fold [11]. Since the Rossman fold is extremely widely distributed in all kingdoms of life, why this NRF-MTase fold has appeared or been conserved through evolution is puzzling. Second, the ExoN activity of nsp14 is a marker of large RNA genomes [21]. The ExoN domain confers capability of RNA mismatch repair to the replicative complex, ensuring genetic stability [11,20,22].

There is still a significant gap of general knowledge on *Nidovirales* RNA capping. For example, *i*) the guanylyltransferase step is still elusive in the whole order; *ii*) if any cap exists in *Arteriviridae*, how is RNA cap methylation achieved ? and *iii*) how is the cap N7-guanine methylated in *Tobaniviridae* and *Roniviridae*, since there is no N7-guanine MTase clearly identified in these families?

In this paper, we report the discovery of an RNA-cap N7-guanine RF-MTase sequence in the Orf1a of ten members of the *Tobaniviridae* family. Remarkably, within this family, this MTase is absent from mammalian toroviruses, but may well represent a genetic marker of non-mammalian *Tobaniviridae*.

## Material and Methods

Virus genome sequences were retrieved from the NCBI database (https://www.ncbi.nlm.nih.gov/genomes/GenomesGroup.cgi?taxid=439488) or Genbank (ncbi.nlm.nih.gov/genbank). The data set (given in Table 1) was that of Lauber *et al*. [21], to which novel or recently described genome sequences (retrieved in NCBI of GenBank) were added purposely. Additionally, the putative and unpublished Palaemon nidovirus and Western flower thrips nidovirus, and the proposed Botrylloides leachii nidovirus [23], were identified in the Sequence Read Archive (SRA) runs SRR5658389, SRR492945 and SRR2729873, respectively, and the virus sequences were assembled, curated and annotated as described elsewhere [24,25]. The resulting sequences are available upon request to H.D.

The conserved Nidovirales C-terminus RdRp core was used to build phylogenetic trees based on MAFFT v7.427 multiple sequence alignment with BLOSUM62 scoring matrix and G-INS-i iterative refinement method. The alignments were used as input for maximum likelihood trees generated with the FasTtree v2.1.5 software (best-fit model = JTT-Jones-Taylor-Thorton with single rate of evolution for each site = CAT). Local support values were computed using the Shimodaira-Hasegawa test (SH) with 1,000 replicates.

Structural alignment of reference or retrieved Mtases were done using EXPRESSO [26] and proofed with Chimera [27]. N7-Mtases reference motifs were inferred and defined based on visual inspection and analysis of the structures. The subsequent motifs were used as orthogonal validation and not used for the search.

Sequences were analyzed using HHblits and HHpred tools of the BioInformatics tookit ([28]). When available, authentic cleavage sites were used to predict protein gene products of the Orf1ab, Orf1a, and Orf1b polyproteins. The boundaries were otherwise approximately (+/-10 aa) determined using structural homologies detected using HHPred, except for the N-term boundary of the Orf1b gene product: In *Nidovirales*, the absence of any structural data nor homology (outside the order) on the N-terminus of the RdRp gene (nsp9 *Arteriviridae*, nsp12 in *Coronaviridae*), which was used for phylogenic analysis, precludes precise sequence homology search in this limited area comprised between the nsp10 and nsp12 proteins (Coronavirus gene-product naming).

Multiple sequence alignments (MSA) were generated using Muscle in SeaView [29]. For each sequence of unknown structure, secondary structures were predicted using Predict Protein [30]. The predicted secondary structures were used to validate the alignment with structural references. The MSA was rendered using ESPript 3.0 [31], together with appropriate structural models as indicated, to assign secondary structures.

When possible, structural 3D models were generated using Phyre 2.0 [32]. Conserved patches of amino-acids were generated using WebLogo [33] and mapped in the structural models rendered in Chimera.

## Results

### All non-arterivirus Nidovirales members carry at least one 2’O-MTase

Available *Nidovirales* genome sequences were used to analyze the presence and primary structure of RNA MTases. Any RF-MTase signature sequence is detectable mainly due to the presence of a typical G-X-G-(X)n-G element part of a SAM-binding motif [34,35], whereas the detection of viral NRF-MTases is based on its homology to the MTase domain of Coronavirus nsp14. We made use of the conserved RdRp fold to build a phylogenetic tree along the whole order. From this tree, we first determined that most of *Nidovirales* code for at least one RF-MTase protein with the canonical K-D-K E catalytic tetrad of 2’O MTase (Fig.1 green & Table 1). Conversely, as previously reported, this RF-2’O MTase is lacking in EAV, LDV, and PRRSV in the *Arteriviridae* family. Interestingly, we identified two RF-MTase signature sequences in the genomes of two recently identified arteri-like viruses : the Hainan Hebius Popei Arterivirus (HHPAV, 12,496 nt) and the Nanhai gost shark arterivirus (NGSAV, 13,162 nt). Their genomes carry a *bona fide* RF-2’-O-MTase signature sequence at the end of their Orf1b, in the gene order RdRp-Hel-(ExoN*)-EndoU-2’OMTase. The NGSAV does not carry a detectable ExoN signature sequence, whereas the HHPAV genome carries a nsp14-like ExoN domain coding region but not a detectable C-terminus NRF-MTase domain, hence the ExoN* labeling (Ferron, Decroly & Canard, unpublished). All these RF-MTase with the K-D-K-E signature are localized in a conserved genomic position at the 3’ end of the Orf1ab, much like their CoV and nsp16 homologues. We performed a MSA of nsp16 from *Roniviridae* and *Tobaniviridae* (Fig. 1, in green), followed by modeling a typical representative of these nsp16 (not shown). All these nsp16-like enzymes appear to be predicted as canonical RNA 2’-O MTases with a typical K-D-K-E tetrad. As noted by others in the Ronivirus nsp16 model [16], small structural differences are observed across the *Tobaniviridae* family, such as the absence of β3 strand and a shorter loop upstream helix αD. Thus, we conclude that non-arterivirus *Nidovirales* code for a RF-MTase with the canonical K-D-K E catalytic tetrad of 2’-O MTase, and that these enzymes are distributed in similar position along genomes.

### The distribution of NRF-MTase is uneven along Nidovirales genomes

As indicated above, many *Nidovirales* members, such as SARS-CoV, stand out because they possess a NRF-MTase, participating to cap N7-guanine methylation, embedded at the C-terminus of their nsp14 gene product. It have been previously reported that the N7-guanine-MTase domain is not uniformly present in nsp14-containing nidoviruses [21]. The phylogenetic tree of Figure 1 reports our findings, and confirms that coronaviruses and most mesoniviruses only possess signature-sequences lining the SAM binding site seen in the nsp14 MTase domain structure. Conversely, the other members of the *Nidovirales* order (*Arteriviridae, Medionivirdae, Roniviridae, Euroniviridae, Abyssoviridae, Mononiviridae* and *Tobaniviridae*) do not seem to embed any nsp14-like NRF-MTase. Two notable exceptions are apparent: First, the unique members of *Abyssoviridae, Mononiviridae* do possess a MTase signature-sequence at the C-terminus of their nsp14-like gene, (i.e., fused to the ExoN domain). Curiously, though, this MTase is readily detectable using HH-Pred as a RF-MTase which does qualify neither as a K-D-K-E 2’-O MTase nor as a N7-guanine MTase (see below). Second, the Fathead Minnow nidovirus 1, belonging to the *Tobaniviridae*, is also an exception as we detected a nsp14-like NRF MTase at the expected genomic Orf1b position. Against, based on the homology with the nsp14-like MTase domain characterized by the absence of K-D-K-E catalytic residue and NRF folding, we hypothesize that these enzymes might play a role in guanine N7 methylation of the RNA cap structure.

Apart from coronaviruses and most mesoniviruses, how nidoviruses methylate their RNA-cap N7-guanine position is thus still unclear.

### What defines N7-guanine MTase features in Nidovirales?

Unlike 2’-O MTases and their well-defined K-D-K-E tetrad, and apart from NRF nsp14-like MTases, signature sequences of RNA cap N7-guanine cap N7-guanosine MTases are much less obvious. The wealth of structural data and mechanistic insight on the N7-guanine MTases enzymes has been acquired from the microsporidian parasite *Encephalitozoon cuniculi* [36], the poxvirus MTase D1:D12 heterodimer [37] and *Reoviridae* [38,39] in the case of RNA viruses. There are only three crystal structure of RF-MTase known to methylate N7-guanine RNA caps of RNA viruses : those of *Reoviridae* (Rotavirus VP4 and Reovirus Lambda2), and Flavivirus, whose NS5 N-terminus domain is a bi-functional N7-guanine and 2’O-MTase [40]. Taking together these data, and combining structural information with the human RNA N7-guanine methyltransferase crystal structure (PDB ID: 3BGV), one cannot define a specific N7-guanine MTase signature (Fig. S1).

From this analysis and others [35] it is clear that the structure conservation prevailed over sequence conservation. No amino acid motifs typical of an RNA N7-guanine MTase can be clearly identified (Fig. S1). A narrower structural comparison including only *Encephalitozoon cuniculi*, poxvirus and the human RNA N7-guanine MTase allows detection of 6 amino acids or motifs K / G/D/ HY / E / Y (Fig. S2). Typical SAM binding motifs are defined by a combination of 3 motifs placed at specific structural positions: K within an α helix, followed by a GxGxG motif within a loop and conserved D. The 3 latter motifs are located in the MTase structural pocket where the N7 methylation supposedly occurs.

In order to use an independent method, we performed a structural search using HHpred on PDB and SCOPe databases [41], using each sequence as a query. This allowed to refine the boundaries of the MTase domain. The structure with the closest hit is a SAM-dependent MTase from *Pectobacterium atrosepticum* and then *Encephalitozoon cuniculi*. A quick superimposition of the retrieved structures with N7-guanine MTase shows that these structures can be structurally aligned but with very limited sequence conservation. Surprisingly, apart from a conserved leucine residue 2 amino acid upstream the poorly conserved SAM binding motif (only the third G is strictly conserved) and a glycine-rich motifs in the fifth beta sheet, nothing else is conserved nor defines an active site (Fig. S3).

Taking these criteria into account, there was no obvious indication of RF N7-guanine MTase features in *Nidovirales* other than the two mentioned above in *Abyssoviridae* and *Mononiviridae*. Currently, based on the detection of RF-MTase fold, one can infer a 2’ O-MTase sequence signature when the K-D-K-E tetrad is present, but not a RNA-cap N7-guanine MTase.

### Selected Tobaniviridae members possess a RF-MTase signature-sequence in Orf1a lacking the canonical 2’O catalytic K-D-K-E tetrad

Then, how do *Roniviridae* and *Tobaniviridae* methylate the RNA-cap N7-guanine position? One possibility would be that these nsp16-like were bi-functional, carrying both 2’-O and N7-guanine methylation activity such as evidenced in Flavivirus NS5 protein, with no obvious signature sequence of this bi-functionality. Another possibility would be that another gene would code for an enzyme performing this methylation, and had escaped detection by standard bioinformatic methods.

We thus performed a larger search of MTase signature-sequences along the whole Orf1ab in all *Nidovirales*: surprisingly, a RF-MTase signature-sequence was detected in Orf1a of 10 members of the *Tobaniviridae* family (Fig. 2). Strictly conserved amino-acids in these new viral MTases define three motifs: 3 glycines of the SAM binding site (G54, 56, and 58 in WBV) 2 residues downstream of a 3 amino acid hydrophobic patches in a β-strand structure, a histidine (H117 immediately followed by either F or Y, and Glu175 (Fig. S4). Interestingly the catalytic K-D-K-E tetrad associated to 2’O-MTase is lacking, suggesting that they could correspond to the missing RNA-cap N7-guanine MTase.

**Figure 2.**
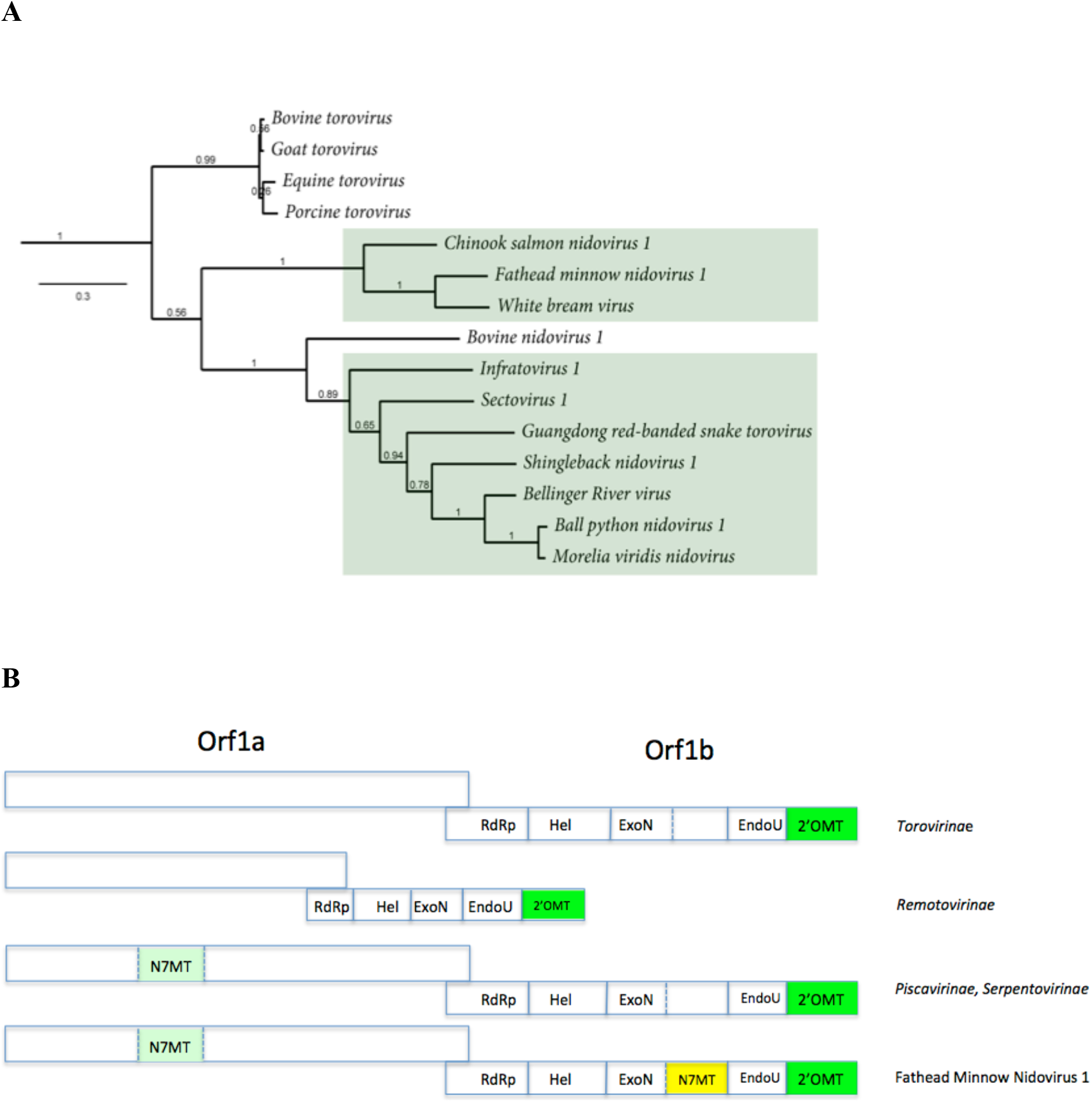
Distribution of the Orf1a MTase along the A) *Tobaniviridae* phylogentic tree (see Fig. 1). B) approximate genome organisation and gene content along the family described in A. The genomes are not drawn to scale. Only the Bostovirus (*Remotovirinae*) genome is represented smaller than its fellow members to account for its ~20 kb genome size.

Remarkably, the novel Orf1a MTase gene was associated with reptiles and fish. Nonetheless, in two cases (Xinzhou Nematode virus 6 and Xinzhou toro-like virus 1, from the *Sectovirus 1* and *Infratovirus 1* species respectively), the viruses were isolated from snake-associated nematodes, although their proper host and life style has not been clearly assessed. The mammalian toroviruses (EToV, BToV, Bovine TCH5 nidovirus, GToV and PToV) do not carry the MTase signature sequence, and thus the latter is associated so far to non-mammalian *Tobaniviridae*.

### The RF-MTase signature-sequence in Orf1a is a putative RNA cap N7-guanine MTase

We performed a structure-based alignment of this Orf1a MTase with the integration of several structurally-defined eukaryotic RNA-cap N7-guanine MTases (Fig. 3A), of which Ecm1 can be considered as prototypic [36]. The conserved amino-acids in these new MTase set remain essentially the same: two glycines of the SAM binding site (G54 and 56 in WBV, G74-76 in Ecm1), a histidine (H117 in WBV, H144 in Ecm1) immediately followed by either F or Y. Glu175 is also very conserved, and in the Ecm1 structure, its homosteric counterpart Glu225 is positioned close to His144 and interacts also with the guanine base, strongly suggesting that this MTase is an N7-guanine MTase. The alignment allows to define 5 conserved motifs which are represented in Fig. 3B, and structurally aligned onto the vaccinia N7-guanine MTase structure D12 (PDB: 4KCB).

**Figure 3.**
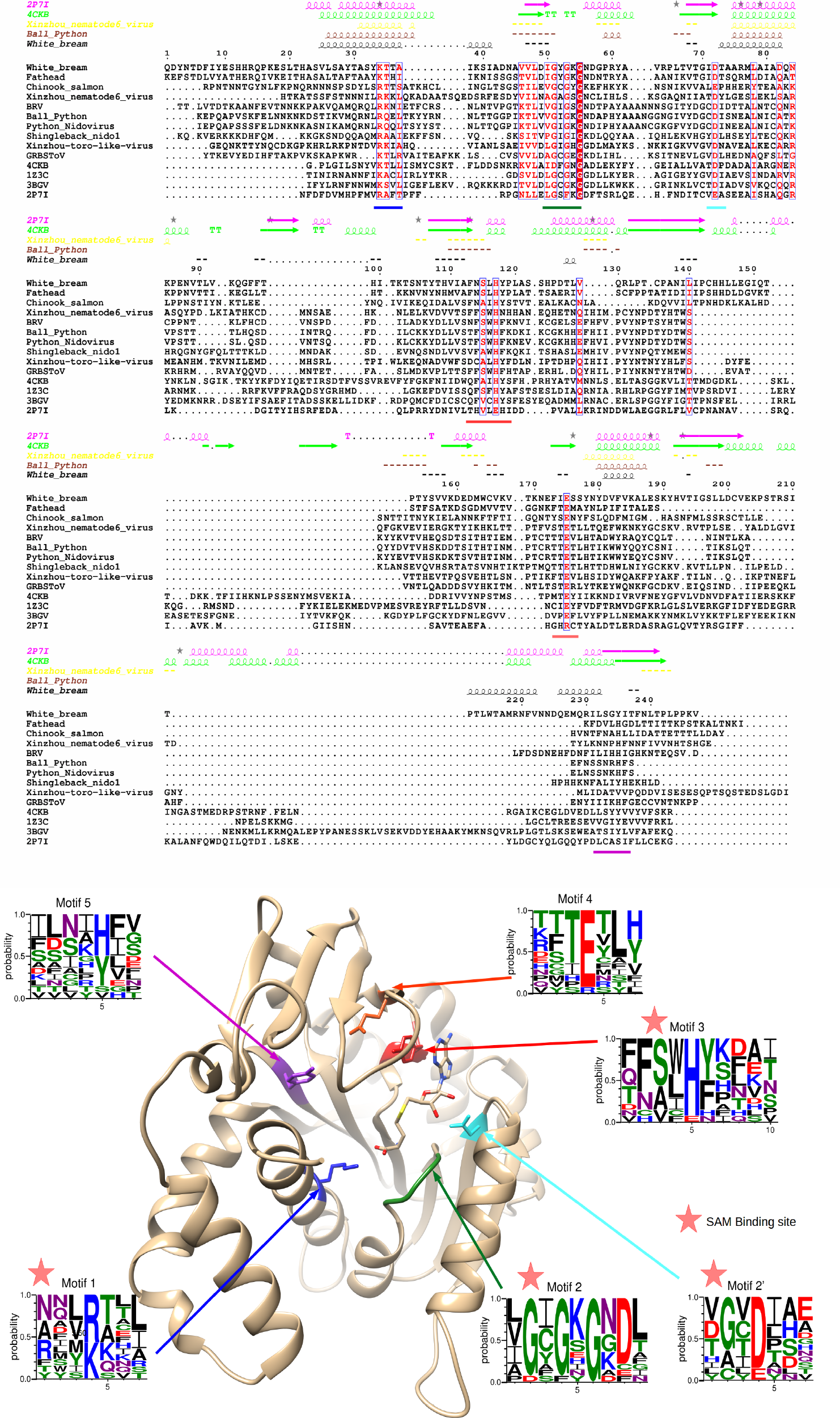
A) Structural alignment of *Tobaniviridae* methyltransferases together with various prokaryotic, eukaryotic and viral members; B) Vaccinia Virus N7-guanine methyltransferases structure (PDB : 4CKB) with highlighted motifs 1 to 5 and their corresponding amino-acid frequencies as determined by WebLogo analysis.

## Discussion

RNA virus cap-MTases perform two types of reactions: the methylation of the N7-guanine of the RNA cap, and the methylation of the adenosine 2’-O ribose of the first transcribed nucleotide [6]. Regarding the guanine methylation, this reaction can also be achieved on a GTP molecule before capping, in some virus families, and the N7-methyl GTP is then used to cap the RNA [42]. This is the case with the alphavirus nsp1 enzyme, whose structure is currently unknown. Conversely, several crystal structures of viral enzymes involved in N7-guanine methylation of RNA caps have been reported, such as that of flaviviruses (NS5, [43]), rotaviruses and reoviruses (VP4 and lambda2, respectively [38,39]), and coronaviruses (nsp14, [11,12]).

The viral 2’-O MTases have been better defined than N7-guanine MTases at the structural and functional level. They all use a K-D-K-E catalytic tetrad, and crystal structures have been determined for several virus groups, such as Reovirus, Flavivirus, Coronavirus, and *Mononegavirales* (reviewed in [6], [44]). All these enzymes (N7-and 2’O–MTase), with SARS-CoV nsp14 as a notable exception (see below), belong to the Rossmann-fold (RF) type MTase (reviewed in [34,35]). They bind the methyl donor (SAM), and are characterized by several structural features: their Rossmann fold is a seven-strands β-sheet surrounded with 6 α-helices; the seventh β-strand is inserted in an anti-parallel orientation between the 5th and 6th strand. The SAM cofactor binds to the first structural motif βαβ, which bears the amino acid sequence motif G-x-G-(x)_n_-G. The RF is an evolutionary ancient fold, which has been widely evolved to perform a variety of chemical reactions. Its structural plasticity is well illustrated by the Flavivirus NS5 MTase, which is able to perform both N7-guanine and 2’-O ribose methylation with the same ~33 kDa domain fused at the N-terminus of the viral RdRp domain [40].

The presence of a MTase in *Nidovirales* Orf1a is unexpected, and had not been reported before, although signature sequences of an uncharacterized MTase had been detected in Ball Python nidovirus, Sectovirus 1 and Infratovirus 1 [23]. Both protein and enzyme functions in *Nidovirales* have been classified into a functional triangle [21], each side of the triangle representing a carrier of the following functions: Orf1a: host defence; Orf1b: genome maintenance; Orfs at the 3’-end: structural products. In this triangle, the MTase-type of activity involved in genome maintenance would usually map to Orf1b [21]. Perhaps indeed this MTase is involved in tasks other than RNA capping, which remain to be determined. Accordingly, it was recently reported that viral or cellular MTases can be recruited by viruses in order to induce internal methylation of their genome [45] and escape to the antiviral response mediated by MDA5 [46]. Out of the gain of this (still putative) enzyme activity in this virus group, the copy number of Orf1a products is expected to be 3-6 time higher than those of Orf1b, raising the following questions: why only these *Tobaniviridae* members need such additional MTase ? Why do they need it in high amounts, provided that the Orf1a stoichiometry is thought to be higher than that of Orf1b products? Are activity levels provided by this enzyme low? What is its actual substrate specificity?

Also, it is legitimate to question the specificity of single copy MTases in *Nidovirales*, such as mammalian *Tobaniviridae* or *Roniviridae* (Fig. 2). Since they do not have any obvious RNA-cap N7-guanine MTase, how is RNA-cap methylation achieved, if any? Are their nsp16-like enzymes of broad or promiscuous specificity?

Our work also uncovers an unexpected 2’-O MTase distribution in viruses seemingly blurring the contour of *Arteriviridae*, a discovery that, in turn, may have a significant impact on our understanding of genome size evolution in *Nidovirales*. Two newly identified arteri-like virus genomes (HHPA and NGSA) provide food for thought regarding RNA genome evolution: Phylogenetic branching of both HHPA and NGSA suggests that arteriviruses might not be primitive small version of larger Coronavirus genomes, but may rather originate from size-reduction of a large Nidovirus ancestor genome.

In any case, the type, diversity, and distribution of RNA MTases along the *Nidovirales* deserves a closer look : SAM-dependent MTases are ancient folds associated with RNA stability and evolution [47]. Their presence and properties in a phylogenetic tree may well give interesting clues regarding RNA genome evolution and its associated issue of host defense mechanisms.

## Acknowledgements

This work was supported by the European Union’s Horizon 2020 research and innovation program through the ANTIVIRALS project under the Marie Sklodowska-Curie grant agreement N° 642434.

## Notes

#### Summary of Updates

An improved definition of the occurence of Non-Rossmann fold MTase has been corrected. Nsp14-like N7-guanine MTase are unique in that they are the only example in the viral world, in addition of being the sole example of Non-Rossmann fold RNA-Cap0 Mtases. This has been corrected throughout the ms.

