## Supplementary material for "Identification of a *Nidovirales* Orf1a N7-guanine cap Methyltransferase signature-sequence as a genetic marker of large genome *Tobaniviridae*": suppl files

### Supplementary Figures

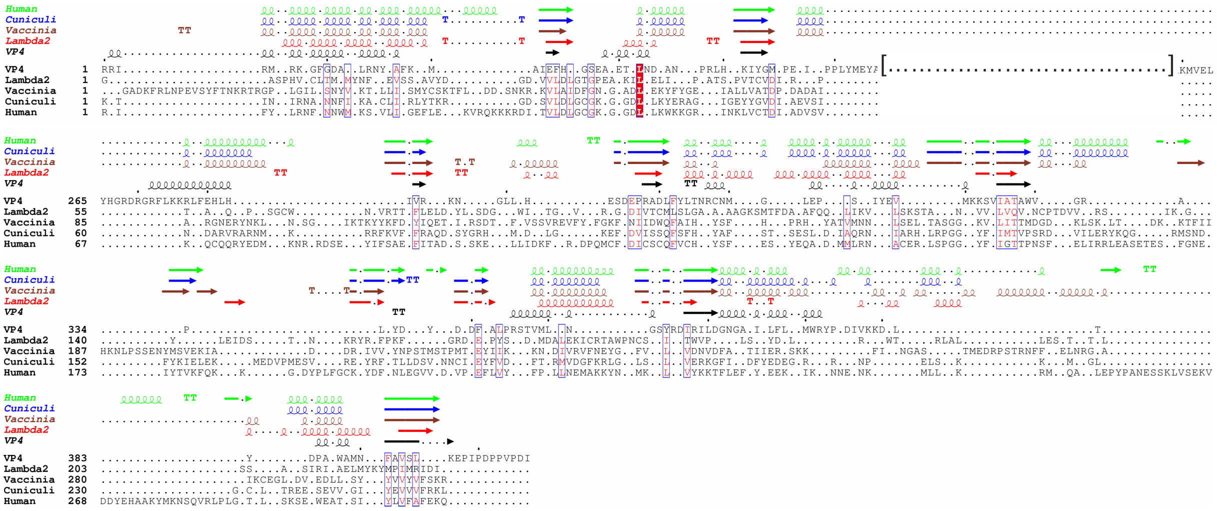

FigS1: Structural alignment of Human methyltransferase (PDB 5E8J [48]), microsporidian parasite *Encephalitozoon cuniculi* (PDB 1RI3 [36]), the poxvirus methyltransferase D1-D12 (PDB 2VDW [37]), and *Reoviridae* in the case of RNA viruses (Rotavirus VP4 (PDB 2JHA [38]) and Reovirus Lambda2 (PDB 1EJ6 [49]).

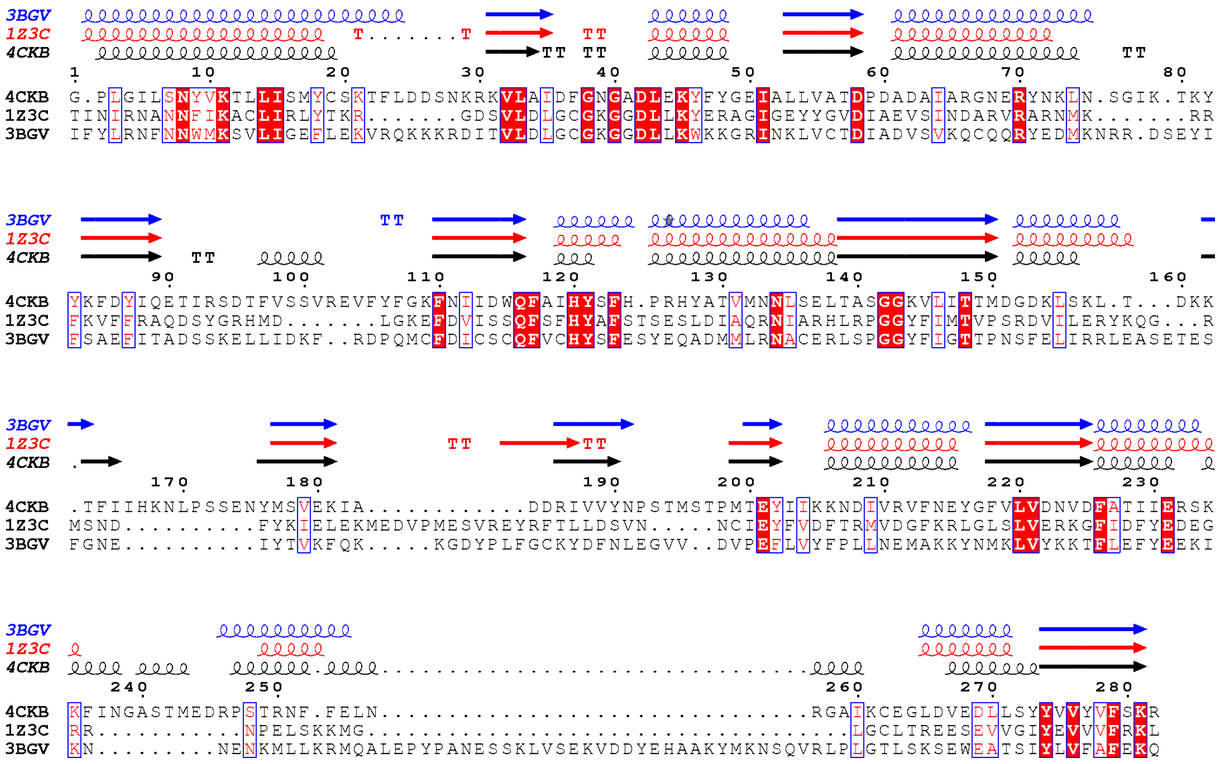


FigS2: Structural alignment of N7 Human methyltransferase (PDB 3BGV ), microsporidian parasite *Encephalitozoon cuniculi* (PDB 1Z3C PMID: [50]), the poxvirus methyltransferase D1-D12 (PDB 4CKB [51]).

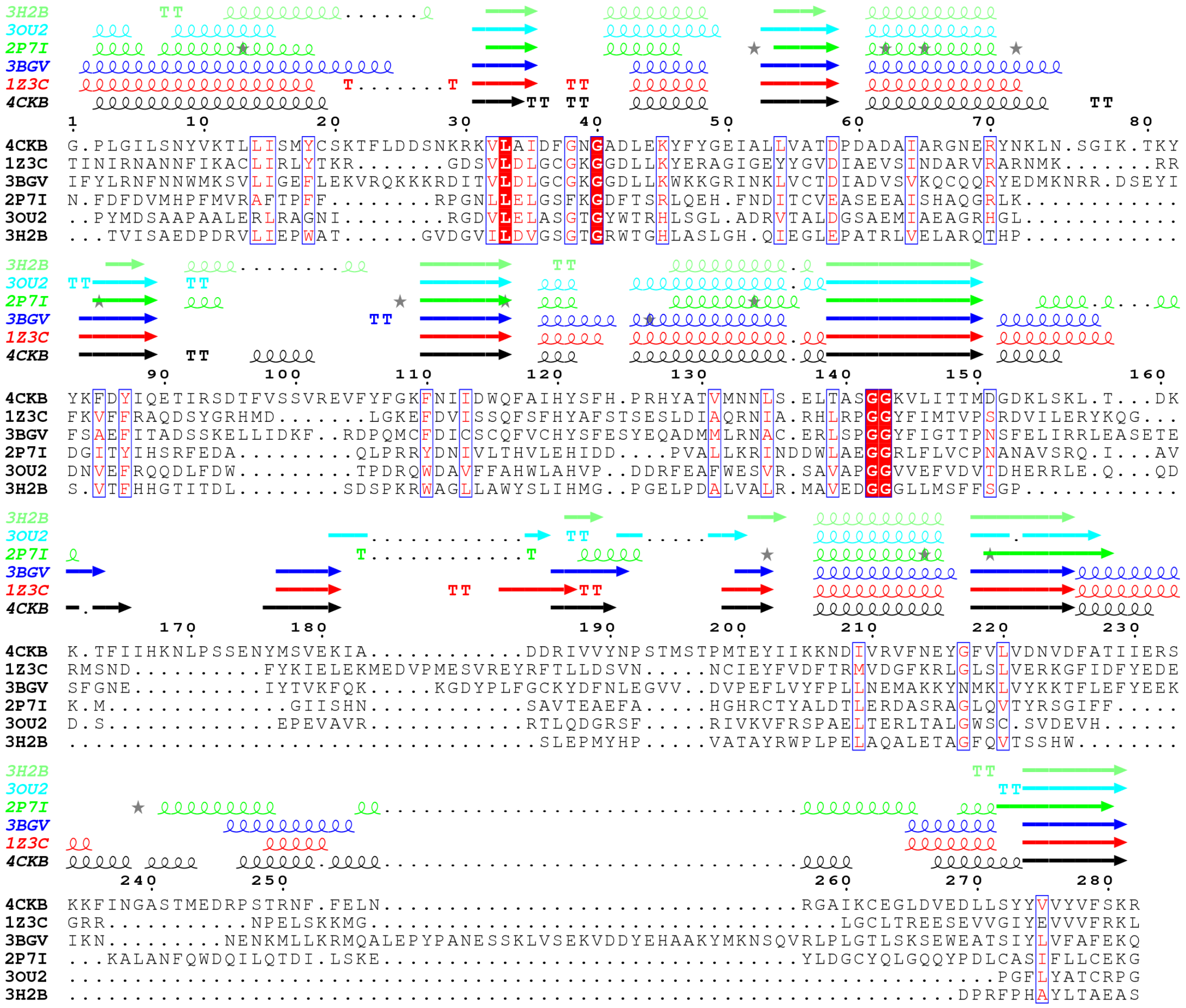


Fig S3: Structural alignment of most common structural hits from HHPRED searches with Nidovirales domain as query : SAM dependent methyl-transferase type 12 family protein (ECA1738) from *Pectobacterium atrosepticum* SCRI1043 (PDB 2P7I), *Streptomyces luridus* methyltransferase *(PDB 3OU2)* and SAM-dependent methyltransferase cg3271 from *Corynebacterium glutamicum (PDB 3H2B) ;* with N7 Human methyltransferase (PDB 3BGV), microsporidian parasite *Encephalitozoon cuniculi* (PDB 1Z3C), the poxvirus methyltransferase D1-D12 (PDB 4CKB).

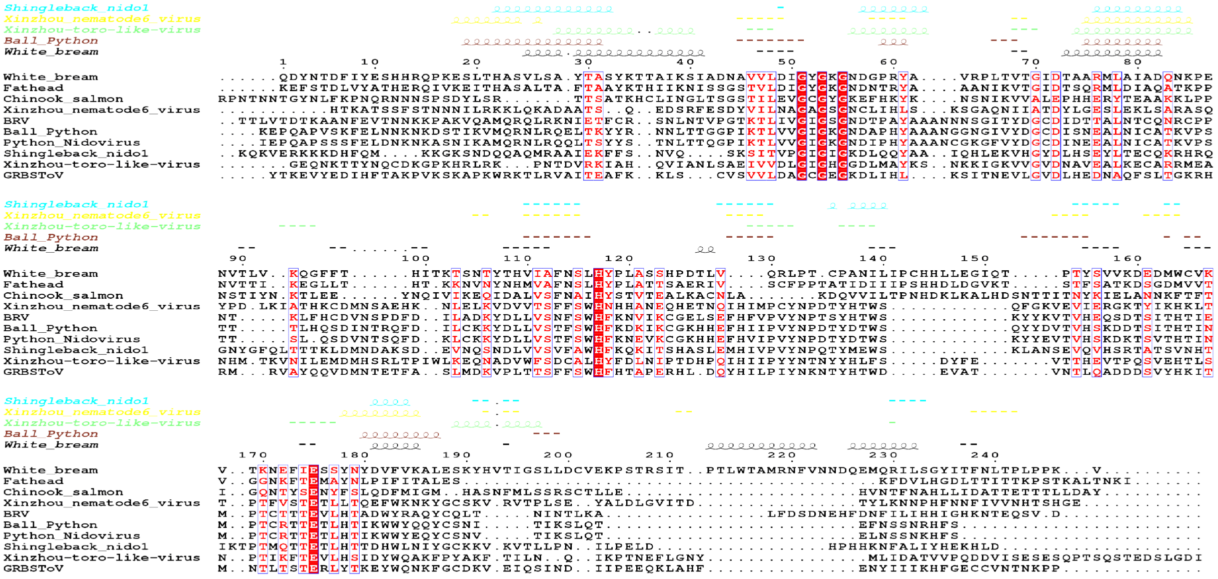

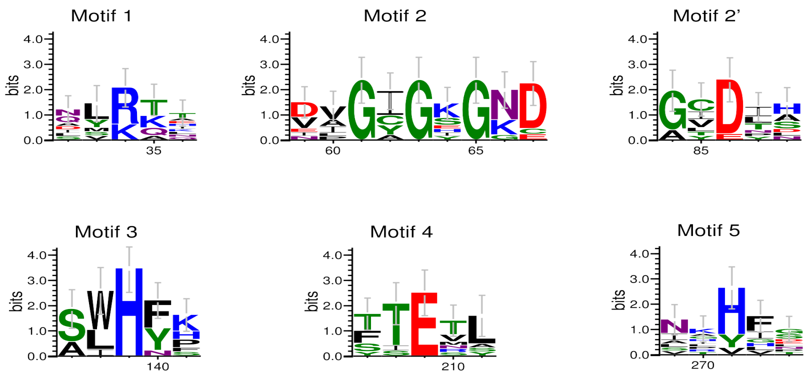


FigS4 : Sequence alignment topped with secondary structure predictions of predicted Methyltransferase domain in *Nidovirales* Orf1a, retrieving 6 motifs : [RK], (GxGxG), D, H[YF], E, [HYv].
